# α4-containing GABA_A_ receptors on dopamine D2 receptor-expressing neurons mediate instrumental responding for conditioned reinforcers, and its potentiation by cocaine

**DOI:** 10.1101/055822

**Authors:** Tom Macpherson, Claire I Dixon, Patricia H. Janak, Delia Belelli, Jeremy J. Lambert, David N. Stephens, Sarah King

## Abstract

Extrasynaptic GABA_A_ receptors (GABA_A_Rs) composed of α4, β and δ subunits mediate GABAergic tonic inhibition and are pertinent molecular targets in the modulation of behavioural responses to drugs of abuse, including ethanol and cocaine. These GABAARs are highly expressed within the nucleus accumbens (NAc) where they influence the excitability of the medium spiny neurons (MSNs). Here we explore their role in modulating behavioural responses to reward-conditioned cues and the behaviour-potentiating effects of cocaine. α4-subunit constitutive knockout mice (α4-/-) showed higher rates of instrumental responding for reward-paired stimuli in a test of conditioned reinforcement (CRf). A similar effect was seen following viral knockdown of GABA_A_R α4 subunits within the NAc. Local infusion of the δ-GABA_A_R-preferring agonist, THIP, into the NAc had no effect on responding when given alone, but reduced cocaine potentiation of responding for conditioned reinforcers in wildtype but not α4-/- mice. Finally, specific deletion of α4-subunits from dopamine D2-, but not D1-receptor-expressing neurons, mimicked the phenotype of the constitutive knockout, potentiating CRf responding and blocking intra-accumbal THIP attenuation of cocaine-potentiated CRf responding. These data demonstrate that α4-GABA_A_R mediated inhibition of dopamine D2 receptor-expressing neurons reduces instrumental-responding for a conditioned reinforcer, and its potentiation by cocaine, and emphasise the potential importance of GABAergic signalling within the NAc in mediating cocaine’s effects.

## Introduction

γ-aminobutyric acid A receptors (GABA_A_RS) are a family of heteropentameric cys-loop ligand-gated chloride channels which function as the major mediators of inhibition in the mammalian CNS. Over 95% of the neurons in the striatum are GABAergic and as such, these receptors are likely to play an important role in modulating the signalling of basal ganglia reward pathways, and in particular the medium spiny neurons (MSNs) of the nucleus accumbens (NAc). The NAc is known to mediate attribution of salience to competing motivational inputs by integrating glutamatergic and dopaminergic inputs (Robbins and Everitt, 1996; Everitt and Robbins, 2005; Nicola, 2006) but little attention is paid to the involvement of GABAergic signalling in these processes. The NAc is also critically involved in Pavlovian associative processes, including conditioned reinforcement (CRf), whereby an environmental conditioned stimulus (CS), which previously held no intrinsic value, acts to support instrumental behaviour following its association with a biologically significant unconditioned stimulus (US) and thereby acquires motivational value (Everitt et al., 2001).

Many drugs of abuse, including cocaine, increase dopamine transmission within the NAc, and potentiate reward-seeking behaviours such as CRf (Taylor and Robbins, 1986).

Synaptic α2-GABA_A_Rs mediate phasic inhibition within the NAc and influence locomotor sensitisation to cocaine, as well as cocaine-potentiation of CRf responding (Morris et al., 2008; Dixon et al., 2010). Additionally, extrasynaptic α4βδ-GABA_A_Rs play an important role in the NAc, mediating a tonic inhibitory conductance, which influences the excitability of MSNs (Maguire et al., 2014). These accumbal extrasynaptic receptors influence ethanol consumption (Rewal et al., 2009; Nie et al., 2011; Rewal et al., 2011). Mice carrying mutations of the GABA_A_R β1 subunit also possess a greatly increased tonic conductance, and these mice, too, exhibit a clear preference for ethanol and are more sensitive to ethanol intoxication (Anstee et al., 2013). However, whether ethanol directly interacts with these receptors remains controversial (Anstee et a., 2013; Lovinger & Homanics, 2007). Our recent finding (Maguire et al., 2014) that activation of α4βδ GABA_A_RS using the selective δ-GABA_A_R agonist THIP (4,5,6,7-tetrahydroisoxazolo[5,4-c]pyridin-3-ol, Gaboxadol) attenuates cocaine potentiation of cocaine conditioned place preference (cocaine-CPP) indicates that these receptors may play a broader role in addictive processes. MSNs within the striatum are typically divided into two subpopulations based upon their expression of dopamine receptors, releasable peptides and their axonal projection targets (Gerfen and Young, 1988). Dopamine D1A-receptor (D1R-), dynorphin-, and substance P-expressing striatonigral neurons, and D2-receptor (D2R-) and enkephalin-expressing striatopallidal neurons, form integral parts of the *direct* and *indirect* basal ganglia pathways, respectively (Gerfen, 1992). While it is known that these two subpopulations act in an opposing manner to control movement, their role in mediating reward learning and control of complex behaviours is still poorly understood. Recent evidence has shown that D1Rs in the NAc mediate Pavlovian conditioned approach and instrumental responding for natural-rewards (Gore and Zweifel, 2013). Alternatively, inhibition of D2R- expressing NAc MSNs using a chemicogenetic approach, or activation using optogenetics, enhances or suppresses motivation to seek cocaine, respectively, suggesting that this MSN population may serve as a protective mechanism against the development of addictive behaviours (Bock et al., 2013). Recently, we demonstrated that α4-GABA_A_Rs on either D1R- or D2R-expressing neurons differentially modulate acquisition of cocaine-CPP and enhancement of its expression by cocaine administration (Maguire et al., 2014), highlighting the possibility that GABAergic inhibition of distinct MSN populations may also regulate other addiction-associated behaviours, including CRf.

Here we test the hypothesis that tonic inhibition through α4-GABA_A_Rs in the NAc influence behavioural responses to natural reward-paired stimuli through a combination of pharmacological activation and deletion of α4-GABA_A_Rs using constitutive GABA_A_R α4-subunit knockout, and α4-subunit viral knockdown mice. Secondly, using cell-specific GABA_A_R α4-subunit knockout mice we explore whether these response are mediated through specific action of α4-GABA_A_Rs on either D1- or D2R-containing neurons. Local infusion of THIP into the NAc in conjunction with cocaine administration reveals a substantial role for α4βδ GABA_A_RS on D2R-expressing neurons in modulating cocaine potentiation of instrumental responding to conditioned reinforcers. These data suggest different roles for α4-GABA_A_Rs on NAc D1R- and D2R-neurons in mediating behaviours associated with addiction to cocaine.

## Materials and Methods

*Production of mice.* Constitutive α4-subunit knockout mice were produced at Sussex University by crossing “floxed” α4 mice (B6.129-*Gabra4^tm1.2Geh^*/J (Jackson Laboratory)), with loxP sites flanking the first coding exon (exon3) of the gabra4 gene, with Cre-recombinase-expressing transgenic mice (see descriptions in (Chandra et al., 2006 and Maguire et al., 2014). Heterozygote mice at Sussex, are maintained on a C57BL/6J background and experimental mice (wildtype (WT) and constitutive α4-subunit knockout mice (α4^-/-^)) generated from subsequent heterozygote matings. Conditional α4 knockout mice with the deletion localized to D1R- or D2R-containing neurons respectively (α4^D1-/-^ or α4^D2-/-^) were created by crossing the “floxed” α4 mice with BAC transgenic mice expressing Cre-recombinase under the control of either the dopamine receptor D1A or D2 gene (B6.FVB(Cg)-Tg(Drd1a-cre)EY266Gsat/Mmucd and B6.FVB(Cg)-Tg(Drd2-cre)ER44Gsat/Mmucd respectively (MMRRC) as described in Maguire et al. (2014). Experimental mice were created from matings of mice both homozygote for the floxed α4 allele, but with one carrying the CRE transgene (cell specific knockout), and one not (wildtype). All mice were on a C57BL/6J background. C57BL/6J male mice (6-8 weeks) were purchased from Charles River, for the viral knockdown studies.

*Animals*. Experimental mice, weighing between 20-30g, were housed in groups of 2-3. Body weights were maintained at approximately 85% of free-feeding weight by the provision of a limited amount of standard lab chow (B&K Feeds, Hull, UK) approximately 2 h after completion of daily experiments. Water was available *ad libitum*. A 12 h light/dark cycle was used (lights on at 7:00 A.M.) with the holding room temperature maintained at 21 ± 2°C and humidity 50 ± 5%. All injections, infusions and behavioural testing were performed between 2:00 P.M. and 5:00 P.M. All procedures were conducted in accordance to the UK Animals (Scientific Procedures) Act 1986, following ethical review by the University of Sussex Ethical Review Committee.

*Stereotaxic Cannulation*. Mice anesthetized with isoflurane were implanted stereotaxically with bilateral guide cannulae (26 ga.) aimed at NAc (coordinates AP1.34; L+/– 1.00; DV −3.20). Following surgery mice were singly housed and allowed to recover for 7 days. A steel infuser (33ga) connected via polyvinyl tubing to a (5µl) Hamilton Gastight syringe was used to infuse 1µl (0.5µl per side) of either saline or THIP bilaterally 1mm beyond the guide cannulae, across 90 seconds, then left to settle for 90 seconds before infusers were removed. The location of cannulae was subsequently confirmed histologically.

*Stereotaxic Viral Infusion*. Mice anesthetized with isoflurane were stereotaxically infused with adenoviruses carrying a shRNA designed to knockdown α4 (Ad-shα4) or a scrambled sequence (Ad-NSS) (Rewal et al., 2009), bilaterally into the NAc (coordinates AP1.34; L+/– 1.40; DV −4.20). A steel infuser (33ga) connected via polyvinyl tubing to a (5µl) Hamilton Gastight syringe was used to infuse 1ul (0.5µl per side) of virus at a rate of 0.2µl/min for 5 minutes, then left to settle for an additional 5 minutes. Following surgery mice were singly housed and allowed to recover for 7 days. The location of infusion was later confirmed using immunofluorescence.

*Conditioned Reinforcement*. Following food deprivation to maintain approximately 85% of baseline body weight, animals were trained in mouse operant chambers (Med Associates Inc, VT, USA), each housed within a light-resistant, sound-attenuating cubicle. Mice underwent 10 consecutive daily 60 minute Pavlovian training sessions during which they were presented with two stimuli, a tone (2.9 KHz, 5 dB above background) or two LED flashing lights. Each stimulus was present 16 times during the session. The order of stimulus presentations was randomly determined and each stimulus trial was separated by a variable, no stimulus, intertrial interval (range of 80-120 seconds; mean = 100 seconds). The stimuli were counterbalanced between mice: one associated with food delivery (CS+), and the other with no outcome (CS−). A single food pellet delivery occurred 5 seconds after CS+ onset (20mg sweetened pellets; 5’TUL, Cat no. 1811142; Test Diets, IN, USA).

Subsequently, two nose-poke detectors were added to the operant chamber, each triggering presentation of either the CS+ or the CS−, (counterbalanced between stimuli). Rates of nose-poke responses were recorded following i.p. cocaine (0, 3, 10, 30 mg/kg) in wildtype and α4^-/-^ mice. Similarly baseline and cocaine- (10mg/kg) potentiated CRf responding were assessed in adenovirus α4 knockdown and control virus-infused mice. Finally, cannulated wildtype, α4^-/-^, and conditional α4^D1-/-^ and α4^D2-/-^ mice, underwent four test days in a latin square design, during which they were administered intra-accumbal infusions of either saline or THIP (3mM) 20 minutes prior to an i.p. injection of saline or cocaine (10mg/kg), directly prior to testing.

*Quantitative reverse transcription PCR.* RNA was extracted from biopsy punches (1mm × 1mm) of the NAc and dorsal striatum (RNeasy Mini Kit, QIAGEN, Limburg, Netherlands) and the amount of RNA was determined using a NanoDrop 2000 micro-volume spectrophotometer (Thermo Fisher Scientific, MA, USA). RNA (250ng) was subjected to reverse transcriptase in the presence of oligo(dT) and 1µl of the resulting cDNA (15µl) was used in a qPCR reaction (SYBRGreen, Sigma-Aldrich, MO, USA) to amplify Gabrα4 (CCACCCTAAGCATCAGTGC *and* CTGAATGGACCA AGGCATTT; and Gapdh (TGTCTCCTGCGACTTCAAC and AGCCGTATTCATTGTCATACC) as a reference gene. For each sample (run in triplicate) the cycle threshold was determined, dCT change from reference gene (gapdh), ddCt (change from Ad-NSS), and the fold change of expression from Ad-NSS treated mice were calculated and analysed according to the method of Pfaffl (Pfaffl 2001).

*Immunofluorescence*. Microtome cut sections (60 µm) of fixed tissue were subjected to immunohistochemistry (as described (Kharazia et al., 2003)). Slices were incubated in rabbit anti-GFP polyclonal primary antibody (1:10,000, Abcam, Cambridge, UK), and then in donkey anti-rabbit secondary antibody (1:300, Jackson ImmunoResearch, PA, USA). Images were acquired using a confocal laser scanning microscope (LSM) (Zeiss, Jena, Germany) and visualized using LSM software (Zeiss, Jena, Germany). *Drugs*. Cocaine hydrochloride was obtained from Macfarlan Smith (Edinburgh, UK). THIP (4,5,6,7-tetrahydroisoxazolo[5,4-c]pyridin-3-ol) was kindly donated by Bjarke Ebert (Lundbeck, Valby, Denmark). Both drugs were dissolved in 0.9% saline, and administered i.p. at an injection volume of 10 ml/kg and intracranially as described above.

*Data Analysis*. qPCR data were analysed using independent-factors analysis of variance (ANOVA) of Gabrα4 mRNA expression relative to Gapdh, with virus treatment as the between-subjects factor. Pavlovian conditioning was assessed using a four-way mixed-factors ANOVA, with genotype as the between-subjects factor, conditioned stimulus (CS+ or CS−) and session as the within-subjects factors, and magazine entries made during presentation of the conditioned stimulus as the dependent variable. The dose-response of cocaine-potentiation of CRf data was analysed using a four-way mixed-factors ANOVA, with genotype as the between-subjects factor, conditioned stimulus and drug dose as the within-subjects factors, and number of nose-poke responses per session as the dependent variable. All other CRf data were analysed using five-way mixed-factors ANOVAs, with genotype as the between-subjects factor, conditioned stimulus, intra-accumbal THIP (or saline) infusions and cocaine (or saline) injections (i.p.) as the within-subjects factors, and nose-poke responses as the dependent variable. *Post hoc* analyses were conducted where appropriate using paired or unpaired t-tests.

## Results

**Increased conditioned reinforcement responding in GABA_a_ R α4^-/-^ mice**. Both wildtype (WT) and constitutive α4-subunit knockout mice (α4^-/-^) were able to learn the reward-predictive properties of the CS+, to a similar extent, as assessed by increased approaches to the food delivery chamber on CS+ presentation (Fig. 1A & 3B). Similarly, both genotypes accurately learned to elicit presentation of the cues via nose-poke responding (Fig. 1B), demonstrating robust conditioned responses. However, in comparison to WTs, α4^-/-^ mice displayed increased instrumental responding for the CS+. Administration of cocaine dose-dependently potentiated instrumental responding for the conditioned reinforcer to the same extent across genotypes, amplifying the initial cocaine-free pattern of responding (Fig. 1C). Neither genotype nor drug had any effect on responding for the CS−.

**Increased conditioned reinforcement responding in α4 viral knockdown mice**. When infused directly into the NAc (Fig 2A), an Ad-shα4 adenovirus, but not a scrambled adenovirus, was able to reduce GABA_A_R α4-subunit mRNA expression (66 ± 6.7% reduction) specifically within the NAc without affecting expression in the neighbouring dorsal striatum (Fig. 2B). As with constitutive knockout mice, both control and α4-subunit viral knockdown mice were able to learn the food-predictive properties of the CS+ (Fig. 2C). Similarly, α4-subunit viral knockdown mice showed an increased instrumental responding for the conditioned reinforcer relative to controls, providing evidence that α4-GABA_A_Rs specifically within the NAc are involved in mediating instrumental responding for reward-conditioned cues (Fig. 2D). Responses for the CS− did not change under any condition.

**Attenuated cocaine-potentiation of conditioned reinforcement responding in THIP-treated α4 wildtype but not α4^-/-^ mice**. Local infusion of THIP (3mM, 210ng per side) into the NAc via indwelling bilateral cannulae (Fig. 3A) did not alter baseline CRf responding. However, THIP was able to decrease cocaine-potentiated responding in WT but not α4^-/-^ mice (Fig. 3C). The specificity of this attenuation to WT mice suggests that selective activation of NAc α4βδ GABA_A_RS by THIP, which does not occur in α4^-/-^ mice (Maguire et al., 2014), modulates responding for conditioned reinforcers. THIP had no effect on responding
for the CS− in either genotype.

**Figure 1.**
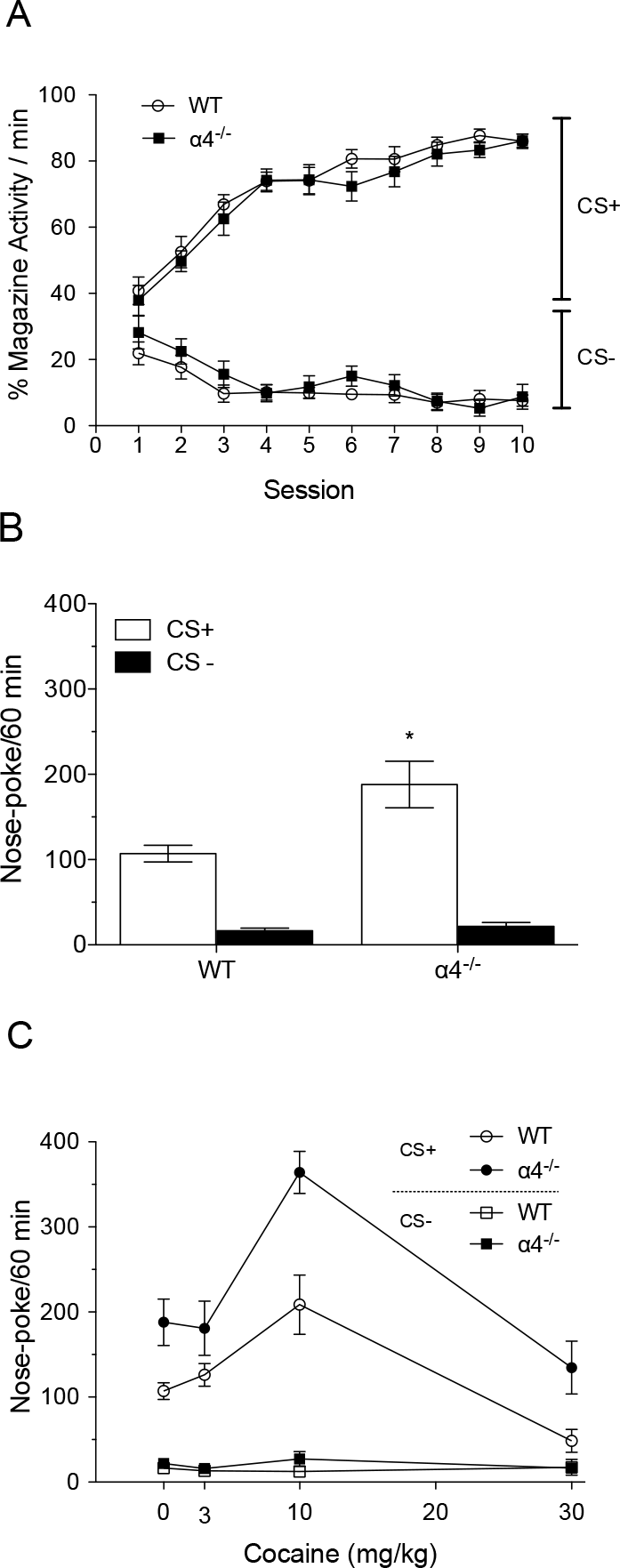
Effects of cocaine on responding in conditioned reinforcement in WT and α4^-/-^ mice. A) In Pavlovian training, WT (n=10) and α4^-/-^ (n=10) mice learnt the association between the cue (CS+) and delivery of a food reward at the same rate (conditioned stimulus by session interaction; F_(9,162)_ = 36.42, p < 0.001). B) In the conditioned reinforcement trial both genotypes preferentially responded on a nose-poke that led to CS+ presentations, compared with a CS− paired nose-poke (main effect of conditioned stimulus, F_(1,18)_ = 334.36, p < 0.001). However, α4^-/-^ mice made significantly more CS+ paired nose-poke responses than WT mice (conditioned stimulus by genotype interaction; F_(1,18)_ = 36.78, p < 0.001). C) Both WT and α4^-/-^ mice showed a cocaine dose-dependent potentiation of responding for the CS+ (conditioned stimulus by drug dose interaction, F_(3,51)_ = 23.97, p < 0.001). This was of similar magnitude in both genotypes. Data are presented as mean +/− SEM. * p < 0.05 compared with WT.

**Figure 2.**
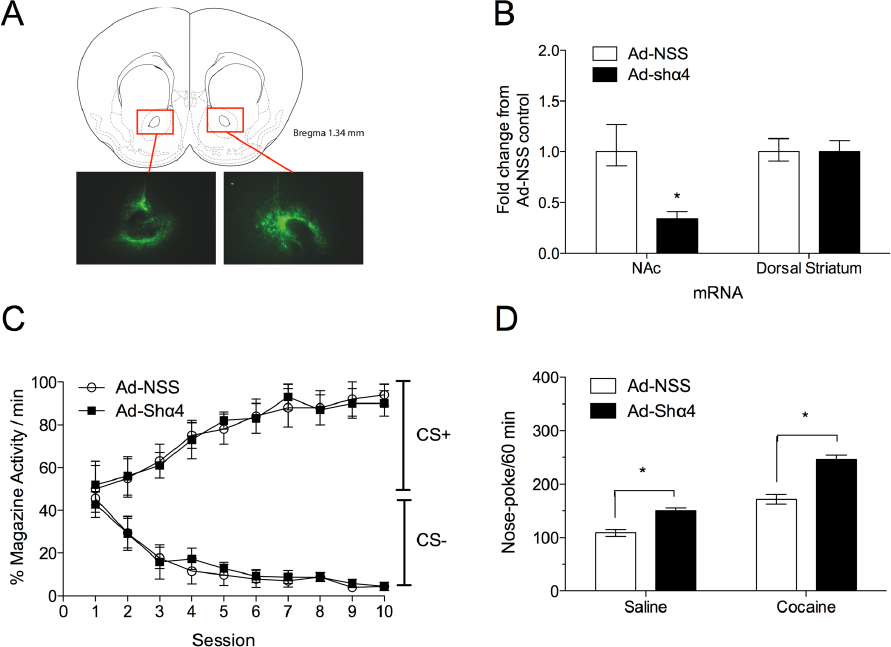
Effects of cocaine on responding in conditioned reinforcement in control and α4-subunit viral knockdown mice. A) Confirmation of virus infusion site in NAc by GFP immunohistochemistry. B) qRT-PCR quantification of mRNA from NAc and dorsal striatum tissue to confirm specificity of Ad-shα4 adenovirus reduction of α4 mRNA expression in the NAc (virus group by brain region interaction, F_(2,15)_ = 32.16, p < 0.001). The histogram depicts fold change expression of Gabra2 from mice infused with control virus (Ad-NSS) ± SEM (n=6 per group). C) Pavlovian training: both control virus (n=10) and Ad-shα4 virus (n=10) infused mice learnt the association between a Pavlovian cue and a food reward at the same rate (conditioned stimulus by session interaction; F_(9,162)_ = 28.13, p < 0.001). D) The Ad-shα4 infused mice showed significantly greater responding for a conditioned reinforcer than Ad-NSS infused mice (conditioned stimulus by virus interaction, F_(1,18)_ = 431.85, p < 0.001). Responding was potentiated by cocaine administration to a similar extent in Ad-NSS and Ad-shα4 infused mice (main effect of drug injection; F_(1,18)_ = 36.57, p < 0.001). Data are presented as mean ± SEM. * p < 0.05 compared with Ad-NSS.

**Targeted deletion of GABA_A_ α4 subunits on D2R-expressing neurons increases responding in conditioned reinforcement and blocks THIP attenuation of cocaine-potentiated responding**. Given that our previous findings have suggested that α4-GABA_A_RS on D1R- or D2R-expressing neurons, respectively, have distinct roles in mediating addiction-associated behaviours (Maguire et al., 2014), we explored the consequences of specifically deleting α4 subunits from these two neuronal populations on instrumental responding for CRf and cocaine-potentiated CRf.

As with the previous experiments, no differences were seen between any of the genotypes in their ability to learn the reward-predictive properties of the CS+ (Fig. 4C & 4D). In a similar manner to constitutive knockouts, D2R-cell specific α4-GABA_A_R knockout mice (α4^D2-/-^) demonstrated increased baseline and cocaine-potentiated CRf responding when compared to their wildtype controls (Fig. 4F). However, this increase was absent in dopamine D1R-cell specific α4-GABA_A_R knockout mice (α4^D1-/-^) (Fig. 4E).

Local infusion of THIP into the NAc did not affect baseline CRf responding in wildtype, or α4^D2-/-^ mice. However, THIP was able to block cocaine-potentiated CRf responding in wildtype and α4^D1-/-^ mice, but not α4^D2-/-^ mice (Fig. 4E & 4F), suggesting that the inability of THIP to block cocaine-potentiated CRf responding in the constitutive α4-knockout mice is due to the absence of α4-GABA_A_RS in NAc D2R- neurons. Responding for the CS− did not change under any condition.

**Figure 3.**
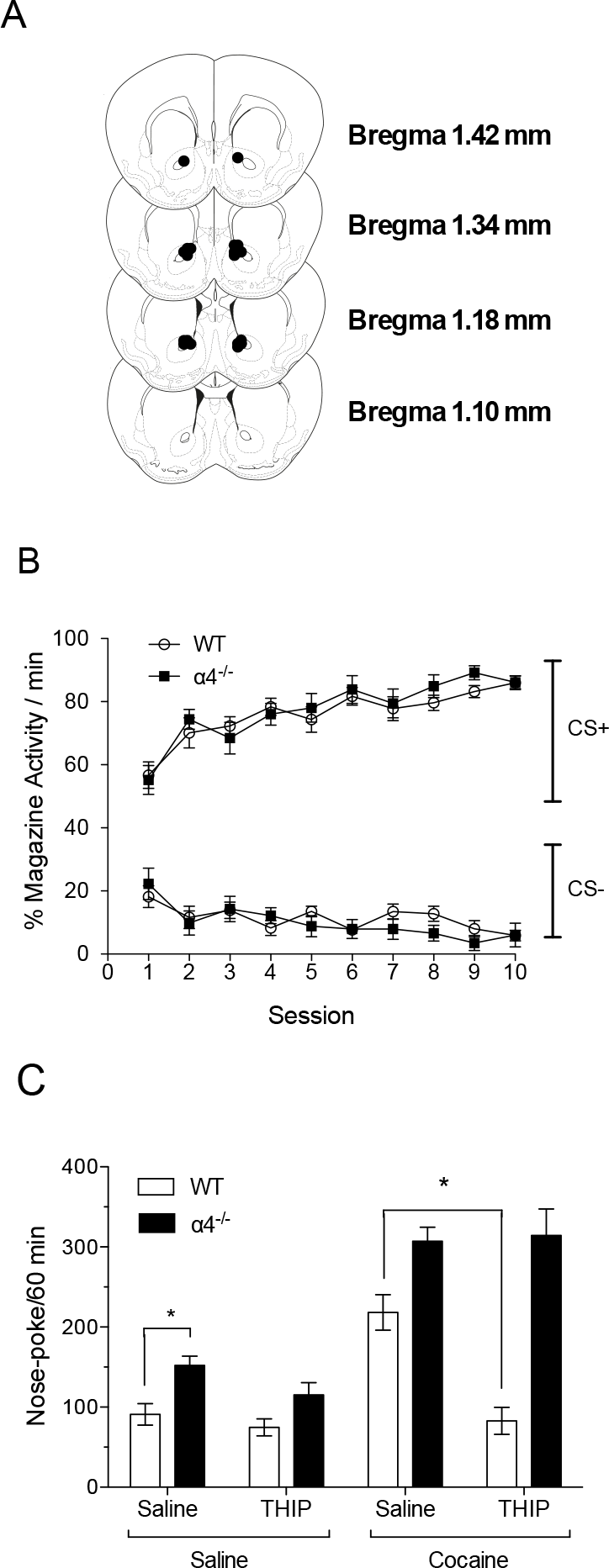
Effects of THIP on responding in conditioned reinforcement in WT and α4^-/-^ mice. A) Cannula placements for intra-accumbal infusions. B) Pavlovian training: WT (n=8) and α4^-/-^ (n=8) mice learnt the association between the cue (CS+) and delivery of a food reward at the same rate (conditioned stimulus by session; F_(9,126)_ = 24.27, p < 0.001). C) Conditioned responding to the CS+ following local NAc infusion of THIP (3mM) and i.p injection of cocaine (10mg/kg). WT but not α4^-/-^ mice displayed an attenuation of cocaine potentiated CS+ responding following intra-NAc THIP infusion (conditioned stimulus by infusion by injection by genotype interaction; F_(1,14)_ = 20.63, p < 0.001). Data are presented as mean ± SEM. *p < 0.05, post hoc paired t test.

**Figure 4.**
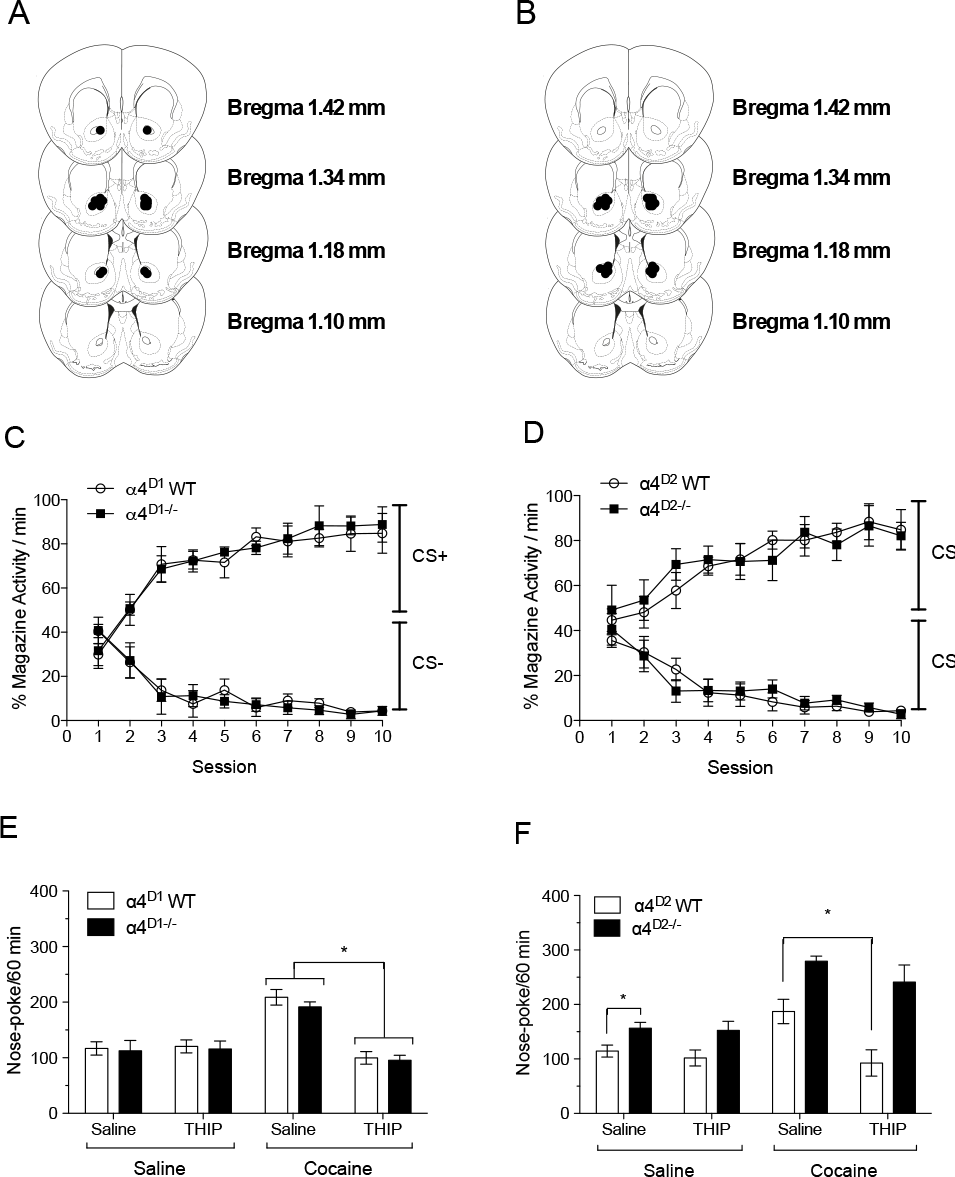
Effect of D1R- or D2R-specific ablation of α4-GABA_A_Rs on conditioned reinforcement with cocaine and THIP. A) Cannula placements for intra-accumbal infusions in WT and α4^D1-/-^ mice. B) Cannula placements for intra-accumbal infusions in WT and α4^D2-/-^ mice. C) Pavlovian training: WT (n=8) and α4^D1-/-^ (n=8) mice learnt the association between the Pavlovian cue and food reward to a similar extent (session by conditioned stimulus interaction, F_(9,279)_ = 70.69, p < 0.001). D) Pavlovian training: WT (n=8) and α4^D2-/-^ (n=8) mice learnt the association between the Pavlovian cue and food reward to a similar extent (session by conditioned stimulus interaction, F_(9,279)_ = 53.98, p < 0.001). E) No difference in CRf responding between WT and α4^D1-/-^ mice, cocaine potentiated responding to a similar magnitude in both genotypes (main effect of drug injection, F_(1,14)_ = 50.14, p < 0.001). THIP alone had no effect on responding but blocked the cocaine potentiation of CS+ responding in both genotypes (conditioned stimulus by infusion by injection interaction; F_(1,14)_ = 10.19, p < 0.01). F) Instrumental responding for the CS+ was increased in α4^D2-/-^ mice compared to WTs (conditioned stimulus by genotype interaction; F_(1,14)_ = 44.38, p < 0.001). Cocaine potentiated responding for the CS+ to a similar degree in both genotypes (main effect of drug injection, F_(1,28)_ = 211.63, p < 0.001). THIP had no effect alone on responding in either genotype, however the attenuation of cocaine potentiation seen in WTs was absent in α4^D2-/-^ mice (conditioned stimulus by infusion by injection by genotype interaction: F_(1,14)_ = 7.70, p < 0.01). Data are presented as mean ± SEM. *p < 0.05, post hoc paired t test.

## Discussion

The data presented here demonstrate that a global deletion of α4-GABA_A_Rs increases instrumental responding for a conditioned reinforcer. Furthermore, although activation of α4-GABA_A_Rs by THIP was not able to reduce baseline CRf responding, it did block cocaine-potentiated responding when infused directly into the NAc, and this effect was abolished in α4^-/-^ mice. The localisation of these α4-GABA_A_Rs actions to the NAc was confirmed by viral knockdown of Gabrα4 expression within NAc producing an increased CRf response comparable to that of the α4^-/-^ mice. Finally, the interaction of α4-GABA_A_R-mediated inhibition with distinct striatal pathways was explored with the use of D1R and D2R-neuron specific GABA_A_R α4-subunit knockout mice. The increased CRf responding in constitutive α4-GABA_A_R knockout and viral knockdown mice was replicated only in D2R-specific α4-GABA_A_R knockout mice. Furthermore the effect of intra-accumbal THIP to reduce cocaine-potentiated CRf responding, was similarly lost in these mice.

The constitutive, cell specific, knockout and viral knockdown mice were able to learn the reward-predictive properties of a conditioned cue to the same extent as their respective controls in all experiments, indicating that α4βδ-GABA_A_Rs play little role in learning the predictive qualities of food-conditioned cues. They also showed similar levels of instrumental responding for a primary reinforcer, food (data not shown), consistent with unchanged responding for sucrose following α4 knockdown in rats (Rewal et al., 2009, 2011). Thus, α4-GABA_A_RS appear not to be involved in Pavlovian associative learning processes, or instrumental responding, but, rather, modulate the expression of behavioural responses specifically to conditioned stimuli.

Activation of dopamine D2Rs reduces α4βδ-GABA_A_R-mediated tonic currents in NAc D2R-MSNs, thereby presumably increasing their excitability (Maguire et al., 2014). Here, suppression of α4-GABA_A_R-mediated inhibition in D2-MSNs causes an increase in responding for a conditioned reinforcer, suggesting that activation of D2-MSNs (possibly indirectly through dopamine receptor activation) potentiates CRf responding. This interpretation is consistent with previous evidence that dopamine D2R agonists quinpirole and bromocriptine both facilitate CRf responding (Beninger et al., 1989; Beninger and Ranaldi, 1992). The current experiments also indicate that the NAc is the site of action of D2R-neuron-mediated potentiation of CRf responding. Mice with a viral knockdown of α4-subunits localized to the NAc demonstrate a similar phenotype to constitutive and D2R-specific α4-GABA_A_R knockout mice, consistent with previous evidence that intra-accumbal administration of quinpirole increases CRf responding (Wolterink et al., 1993).

The potentiation of CRf responding by psychostimulants also appears to be mediated by D2R-MSNs within the NAc. In the current experiments, intra-accumbal THIP was able to block the cocaine-potentiation of CRf responding in wildtype but not their counterpart constitutive or D2R-specific α4-GABA_A_R knockout mice. This role of D2-MSNs is further supported by evidence that intra-accumbal administration of D2-antagonist raclopride blocks amphetamine-induced potentiation of CRf responding (Wolterink et al., 1993).

If an action of dopamine agonism is to reduce tonic GABAergic currents in D2R- neurons (Maguire et al., 2014), then THIP would be expected to oppose this effect as it would override the D2R-mediated decreased tonic inhibition. However, the dopamine-induced decrease of the tonic current was observed only when NAc D2R- MSNs were incubated with dopamine agonists for a prolonged period (Maguire et al., 2014). Thus the reduced inhibition may not be a rapidly induced effect, and may become apparent only after sustained activation of dopamine D2 receptors, as would occur following cocaine administration (Maguire et al., 2014). If under normal physiological conditions dopamine levels are not sufficient to influence the GABA_A_R tonic current then THIP would have nothing to oppose, explaining why THIP has no behavioural effect in the absence of cocaine.

These data also indicate that there are dissociable roles for α4-GABA_A_Rs on D1- or D2-expressing NAc MSNs in mediating various aspects of reward. Previously we have demonstrated that α4-GABA_A_Rs on D1- but not D2-expressing neurons are involved in the expression of cocaine-CPP and its potentiation by cocaine (Maguire et al., 2014). Thus we have uncovered a possible dichotomy between the striatal pathways involved in controlling behavioural responses to discrete (CRf) and contextual (CPP) conditioned cues. An alternative interpretation that the difference between the present report on food-conditioned discrete cues and our previous report on cocaine-conditioned contextual cues relate to differences in the primary reward is unlikely as we found similar effects of constitutive α4-deletion on food CPP (data not shown) as we have reported for cocaine CPP.

In conclusion, our results indicate that α4-GABA_A_R inhibition of D2R-expressing neurons, probably within the NAc, is a critical mechanism for controlling the expression of the reinforcing properties of discrete reward-conditioned cues. These data suggest that accumbal α4βδ GABA_A_RS may provide a potential therapeutic target for the treatment of addictive behaviours.

## Acknowledgements

This work forms part of the PhD thesis of TM, who was supported by an MRC studentship. The study was also supported by the Medical Research UK (G0802715 & G1000008), an MRC Centenary Award to TM, Tenovus and an Anonymous Trust Grant (D.B. and J.J.L) and NIH R01 AA016835 (P.H.J). The work was carried out within the MRC Addiction Cluster “GABA_A_ receptors in neurobiology of drug and alcohol addictions”.

## References

Anstee QM, Knapp S, Maguire EP, Hosie AM, Thomas P, Mortensen M et al (2013) Mutations in the Gabrb1 gene promote alcohol consumption through increased tonic inhibition. Nature Communications 4:2816–2838.

Beninger RJ, Hoffman DC, Mazurski EJ (1989) Receptor subtype-specific dopaminergic agents and conditioned behavior. Neuroscience & Biobehavioral Reviews 13:113–122.

Beninger RJ, Ranaldi R (1992) The effects of amphetamine, apomorphine, SKF 38393, quinpirole and bromoeriptine on responding for conditioned reward in rats. Behav Pharmacol 3:155.

Bock R, Shin JH, Kaplan AR, Dobi A, Markey E, Kramer PF et al (2013) Strengthening the accumbal indirect pathway promotes resilience to compulsive cocaine use. Nature Neuroscience 16:632–638.

Chandra D, Jia F, Liang J, Peng Z, Suryanarayanan A, Werner DF, et al (2006) GABAA receptor 4 subunits mediate extrasynaptic inhibition in thalamus and dentate gyrus and the action of gaboxadol. Proceedings of the National Academy of Sciences USA 103:15230–15235.

Dixon CI, Morris HV, Breen G, Desrivieres S, Jugurnauth S, Steiner RC et al (2010) Cocaine effects on mouse incentive-learning and human addiction are linked to 2 subunit-containing GABAA receptors. Proceedings of the National Academy of Sciences USA 107:2289–2294.

Everitt BJ, Dickinson A, Robbins TW (2001) The neuropsychological basis of addictive behaviour. Brain Res Brain Research Reviews 36:129–138.

Everitt BJ, Robbins TW (2005) Neural systems of reinforcement for drug addiction: from actions to habits to compulsion. Nature Neuroscience 8:1481–1489.

Gerfen CR (1992) The neostriatal mosaic: multiple levels of compartmental organization in the basal ganglia. Annual Review of Neuroscience 15:285–320.

Gerfen CR, Young WS (1988) Distribution of striatonigral and striatopallidal peptidergic neurons in both patch and matrix compartments: an in situ hybridization histochemistry and fluorescent retrograde tracing study. Brain Research 460:161–167.

Gore BB, Zweifel LS (2013)Genetic reconstruction of dopamine d1 receptor signaling in the nucleus accumbens facilitates natural and drug reward responses. The Journal of Neuroscience 33:8640–8649.

Lovinger DM, Homanics GE (2007) Tonic for what ails us? high affinity GABAA receptors and alcohol. Alcohol 41:139–143

Maguire EP, Macpherson T, Swinny J, Dixon CI, Herd MB, Belelli D et al (2014) Tonic inhibition of accumbal spiny neurons by extrasynaptic α4pS GABAA receptors modulates the actions of psychostimulants. The Journal of Neuroscience 34:823–38.

Morris HV, Dawson GR, Reynolds DS, Atack JR, Rosahl TW, Stephens DN (2008) Alpha2-containing GABAA receptors are involved in mediating stimulant effects of cocaine. Pharmacology, Biochemistry and Behavior 90:9–18.

Nicola SM (2006) The nucleus accumbens as part of a basal ganglia action selection circuit. Psychopharmacology 191:521–550.

Nie H, Rewal M, Gill TM, Rona D, Janak PH (2011) Extrasynaptic delta-containing GABA(A) receptors in the nucleus accumbens dorsomedial shell contribute to alcohol intake. Proceedings of the National Academy of Sciences USA 108:4459–4464.

Pfaffl MW (2001) A new mathematical model for relative quantification in real-time RT-PCR. Nucleic Acids Research 29:2002–2007.

Rewal M, Donahue R, Gill TM, Nie H, Ron D, Janak PH (2011) Alphα4 subunit-containing GABAA receptors in the accumbens shell contribute to the reinforcing effects of alcohol. Addiction Biology 17:309–321

Rewal M, Jurd R, Gill TM, He D-Y, Ron D, Janak PH (2009)α4-Containing GABAA Receptors in the Nucleus Accumbens Mediate Moderate Intake of Alcohol. The Journal of Neuroscience 29:543–549.

Robbins TW, Everitt BJ (1996) Neurobehavioural mechanisms of reward and motivation. Current Opinion Neurobiology 6:228–236.

Taylor J, Robbins T (1986) 6-Hydroxydopamine lesions of the nucleus accumbens, but not of the caudate nucleus, attenuate enhanced responding with reward-related stimuli produced by intra-accumbens d-amphetamine. Psychopharmacology 90:390–397.

Wolterink G, Phillips G, Cador M, Donselaar-Wolterink I, Robbins TW, Everitt BJ (1993) Relative roles of ventral striatal D1 and D2 dopamine receptors in responding with conditioned reinforcement. Psychopharmacology 110:355–364.

